# Intermittent cocaine self-administration has sex-specific effects on addiction-like behaviors in rats

**DOI:** 10.1101/2022.12.19.521063

**Authors:** Brooke N. Bender, Mary M. Torregrossa

## Abstract

Intermittent access (IntA) models of cocaine self-administration were developed to better model in rodents how cocaine is used by human drug users. Compared to traditional continuous access (ContA) models, IntA has been shown to enhance several pharmacological and behavioral effects of cocaine, but few studies have examined sex differences in IntA. Moreover, no one has examined the efficacy of cue extinction to reduce cocaine seeking in the IntA model, which has previously shown to be ineffective in other models that promote habit-like cocaine seeking. Therefore, rats were implanted with jugular vein catheters and dorsolateral striatum (DLS) cannulae and trained to self-administer cocaine paired with an audiovisual cue with ContA or IntA. In subsets of rats, we evaluated: the ability of Pavlovian cue extinction to reduce cue-induced drug seeking; motivation for cocaine using a progressive ratio procedure; compulsive cocaine taking by pairing cocaine infusions with footshocks; and dependence of drug-seeking on DLS dopamine (a measure of habit-like behavior) with the dopamine antagonist *cis*-flupenthixol. Overall, cue extinction reduced cue-induced drug seeking after ContA or IntA. Compared to ContA, IntA resulted in increased motivation for cocaine exclusively in females, but IntA facilitated more compulsive cocaine taking exclusively in males. After 10 days of IntA training, but not fewer, drug-seeking was dependent on DLS dopamine most notably in males. Our results suggest that IntA may be valuable for identifying sex differences in the early stages of drug use and provide a foundation for the investigation of the mechanisms involved.

**Highlights:**

- IntA promotes increased motivation for cocaine in females
- IntA augments compulsive cocaine self-administration in males
- IntA promotes DLS dopamine-dependent cocaine seeking, most notably in males
- Cue extinction overall reduces cue-induced drug seeking after ContA or IntA
- Under IntA, females self-administer more cocaine when in estrous

## 1. Introduction

The progression from recreational drug use to substance use disorder (SUD) involves escalated drug intake, increased motivation for the drug, maladaptive learning and memory, and persistence in drug seeking and taking despite negative consequences (Belin-Rauscent et al., 2016; Bender and Torregrossa, 2020; Edwards and Koob, 2013; Koob and Le Moal, 2008; Smith and Laiks, 2017). Rodent cocaine self-administration models are used to evaluate these and many other characteristics of SUDs and the circuits involved in order to inform understanding of SUDs and develop novel treatments (Panlilio and Goldberg, 2007). Recently, intermittent access (IntA) cocaine self-administration has been implemented to more accurately model human patterns of cocaine use, which typically consist of short periods of drug consumption separated by lengthy periods without consumption (Allain et al., 2015; Samaha et al., 2021). Whereas rodents self-administering under traditional continuous access (ContA, either short [ShA] or long access [LgA]) models tend to maintain high, sustained brain cocaine concentrations throughout sessions, IntA models produce the spiking patterns observed in human cocaine use (Allain et al., 2015; Bentzley et al., 2014; Samaha et al., 2021).

The spiking brain-cocaine concentration patterns characteristic of IntA have different pharmacological and behavioral effects than the continuous elevated cocaine levels sustained in ContA models. IntA increases cocaine-induced dopamine release and uptake and promotes sensitization of the dopamine transporter (DAT) to cocaine in the nucleus accumbens, whereas LgA promotes tolerance (Calipari et al., 2015, 2013; Kawa et al., 2019). Several studies have shown that IntA increases motivation for cocaine (Algallal et al., 2020; Calipari et al., 2015; James et al., 2019; Zimmer et al., 2012), and this enhanced motivation is correlated with sensitization of cocaine-induced dopamine release in the nucleus accumbens (NAc) (Kawa et al., 2019). Because dopamine in the NAc is also important for cocaine-cue associations, it is unsurprising that IntA also augments cue-induced reinstatement (Aragona et al., 2009; James et al., 2019; Kawa et al., 2019), but the ability to extinguish cocaine-cue associations after IntA has not been examined. Compulsive cocaine taking, an important aspect of SUDs, is also enhanced by IntA (Belin-Rauscent et al., 2016; James et al., 2019; Lüscher et al., 2020). Although it has been theorized that compulsive drug seeking arises from an over-reliance on habitual behavior compared to goal-directed drug seeking, the effects of IntA on habit formation or reliance on dopamine in the dorsolateral striatum (DLS), a structure important for habit initiation, have not been evaluated (Everitt, 2014; Ostlund and Balleine, 2008). One study showed that compared to LgA, IntA increased cocaine-induced *c-fos* expression in the dorsal striatum, including the DLS, but not the nucleus accumbens, suggesting IntA may impact habit circuitry, but its behavioral effects on habits are unknown (Minogianis and Samaha, 2020).

Additionally, sex differences in the effect of IntA on addiction-like behaviors are understudied, though existing studies show that IntA produces greater enhancement of psychomotor sensitization, locomotor sensitization, and greater and more rapid enhancement of motivation for cocaine in females (Algallal et al., 2020; Carr et al., 2020; Kawa and Robinson, 2019). The ability of IntA to enhance compulsive cocaine self-administration has only been evaluated in males and has not been examined in females (James et al., 2019). Furthermore, a recent study showed that dopamine sensitization, which appears to be central to IntA’s effects on addiction-like behavior, differentially affects the rate of habit formation in males and females (Schoenberg et al., 2022), suggesting that IntA could have differential effects on the rate at which DLS dopamine-dependent, habit-like cocaine seeking is established between sexes.

Overall, IntA has been shown to promote many addiction-like behaviors characteristic of SUDs and provides a model for analyzing how the spiking brain cocaine concentrations that induce sensitization may promote the progression from casual substance use to SUD. We sought to expand on the current literature by examining the effects of IntA on the ability of cue extinction to reduce cue-induced cocaine seeking, motivation for cocaine on a progressive ratio (PR) schedule, compulsive cocaine self-administration, and the development of DLS dopamine-dependent cocaine seeking in both males and females. Our results support previous findings that IntA-induced enhanced motivation for cocaine is more prominent in females, provide new evidence that IntA promotes DLS dopamine-dependent, habit-like behavior, and suggest that IntA may be valuable in examining sex differences in the early phases of the progression toward SUDs.

## 2. Methods

### 2.1 Animals

Adult Sprague-Dawley rats (Envigo) age 8-9 weeks upon arrival (n=120; male n=60; female n=60) were used. Rats were given at least 5 days to acclimate to the colony before experimentation. Before surgery, rats were pair-housed in autoventilated racks with automated watering in a humidity- and temperature-controlled room with a 12-hour light-dark cycle and had *ad libitum* access to food and water. Rats were individually housed after surgery, and one day before the start of self-administration they were food-restricted to a maintain 90% of their free-feeding body weight. Behavioral experiments were run in the dark cycle under red light and began within 3 hours of the same time of day. Procedures were conducted in accordance with the National Institute of Health’s *Guide for the Care and Use of Laboratory Animals* and were approved by the University of Pittsburgh’s Institutional Animal Care and Use Committee.

### 2.2 Drugs

Cocaine hydrochloride (graciously provided by NIDA) was dissolved at 1 mg/ml in 0.9% sterile saline (Thermo Fischer) and filter-sterilized. *Cis*-flupenthixol hydrochloride (Cayman Chemical Company) was dissolved at 20 μg/μl in ddH_2_O.

### 2.3 Behavioral Apparatus

Experiments were conducted in 24 standard operant conditioning chambers (MedAssociates) using MedPC software (MedAssociates) as previously described (Bender and Torregrossa, 2021). Half of the boxes were equipped with grid floor harnesses that allowed an aversive stimulus generator to pass electrical current through the grid floors which were checked prior to testing with an amp meter (MedAssociates).

### 2.4 Surgery

#### 2.4.1 Anesthesia

Rats were fully anesthetized with ketamine (87.5-100 mg/kg, Henry Schein) and xylazine (5 mg/kg, Butler Schein), were administered the analgesic Rimadyl (5 mg/kg, Henry Schein) and 5 ml Lactated Ringer’s solution, and surgical sites were prepped as previously described (Bender and Torregrossa, 2021; Rich et al., 2019).

#### 2.4.2 Intravenous catheterization

Rats were implanted with a chronic indwelling intravenous catheter (made in-house) into the right jugular vein that was capped to prevent blockages as previously described (Bender and Torregrossa, 2021; Torregrossa and Kalivas, 2008).

#### 2.4.3 Intracranial cannulation

Rats used for experiments involving intra-DLS infusions were implanted with bilateral 22 gauge guide cannulae (PlasticsOne) 1mm dorsal to the DLS (in mm from Bregma, anterior and posterior (AP): +0.8; medial and lateral (ML): +/− 3.0; dorsal and ventral (DV): −4.0)) as previously described (Bender and Torregrossa, 2021) immediately following jugular vein catheterization.

#### 2.4.4 Post-operative care

Rats were administered Rimadyl (5 mg/kg) subcutaneously on the two days following surgery, and catheter patency was maintained with a gentamicin (3 mg/ml; Henry Schein) and heparin (30 USP/ml; Henry Schein) mixture in 0.9% sterile saline as previously described (Bender and Torregrossa, 2021).

### 2.5 Behavioral Procedures

#### 2.5.1 Cocaine self-administration

Rats were trained to self-administer cocaine (0.5 mg/kg/infusion) daily. Each session began with the illumination of the house light, start of the fan, and insertion of the active and inactive lever (counterbalanced between animals). All cocaine infusions were paired with a 10-second audiovisual conditioned stimulus (CS) of a tone and the illumination of a cue light above the active lever and initiated a 10-second time-out period when the house light was extinguished and another infusion could not be obtained. Inactive lever presses were recorded, but had no programmed consequences.

Rats underwent initial acquisition training for 2-10 days, during which cocaine was delivered on a fixed-ratio 1 (FR1) schedule continuously for 125 minutes or until 60 infusions were obtained to reduce overdose risk. Rats met acquisition criteria when they received ≥15 infusions with twice as many active vs. inactive lever presses for two consecutive days, as adapted from previous experiments, adjusting for dose of cocaine (Allain and Samaha, 2019).

After meeting acquisition criteria, rats continued self-administering with either continuous access (ContA) or intermittent access (IntA) to cocaine. IntA sessions consisted of 5 5-minute cocaine-available periods separated by 25 minutes of cocaine-unavailable periods for a total of 125 minutes, a protocol adapted from previous IntA experiments (Allain and Samaha, 2019; Garcia et al., 2020). We used this shortened, 125-minute daily self-administration IntA model because evidence suggests shortened IntA produces similar effects to the original 6-hour IntA paradigm (Allain and Samaha, 2019), and this shortened procedure is more readily applied within labs with limited resources. During cocaine-unavailable periods, the house light was extinguished and levers were retraced, but the fan remained on. The number of total infusions allowed was not restricted, since timeouts and cocaine unavailable periods reduce risk of overdose. In ContA sessions, cocaine was available continuously at the start of the session until a maximum of 30 infusions was reached, after which the house light was extinguished and levers were retraced.

#### 2.5.2 Pavlovian cue extinction

Rats in experiment 1 (n=72; male n=36; female n=36) underwent Pavlovian cue extinction or a control procedure on the day following the tenth day of ContA or IntA. During cue extinction or a 0-CS control procedure, rats were presented with 0 or 120 audiovisual cues noncontingently in the context where self-administration previously occurred with levers retracted. Ten-second cues (previously paired with cocaine infusions) were presented every 30 seconds for 1 hour.

#### 2.5.3 Cue-induced cocaine-seeking test

Rats in experiment 1 underwent a 1-hour cue-induced cocaine-seeking test on the day following cue extinction. During this test, lever presses resulted in cue presentations and timeouts as previously occurred during self-administration, but no cocaine was delivered.

#### 2.5.4 Compulsive cocaine self-administration testing

A subset of rats from experiment 1 (n=24; male n=12; female n=12) underwent testing for compulsive cocaine self-administration after the cue-induced cocaine-seeking test. For 3 days, all rats self-administered cocaine with continuous access in 110-minute sessions. These 3 days allowed IntA-trained rats to learn to revert their cocaine intake back to a load-and-maintain pattern, which prevents binge-like intake from interfering with results of the subsequent compulsion testing (Bentzley et al., 2014; James et al., 2019).

The following day, rats underwent compulsion testing using a protocol previously described (Bentzley et al., 2014; James et al., 2019). Cocaine self-administration was separated into 10-minute bins. During the first bin, rats reached their desired brain cocaine concentration, and during the second bin they were allowed to maintain this unpunished. For the remaining 9 bins, cocaine infusions (0.5 mg/kg/inf) were paired with a 0.5-second aversive footshock delivered at the time of cocaine infusion, and the intensity of this footshock increased incrementally on a tenth-log_10_ scale during each bin, starting at 0.13 mA and ending at 0.79 mA. The charge withstood during each bin was calculated by multiplying the infusion number with the duration of the shock (0.5 s) and the shock amplitude as previously described, and maximum shock withstood in any one bin represents the maximum charge each rat is willing to withstand to maintain their desired brain cocaine concentration, which controls for individual differences in brain cocaine concentration preference (Bentzley et al., 2014).

#### 2.5.5 Progressive ratio

A subset of rats from experiment 1 (n=48; male n=24; female n=24) underwent progressive ratio (PR) testing after the cue-induced cocaine-seeking test. Rats were allowed to self-administer cocaine for 1 day on their previous access model, then underwent a PR protocol as previously described (Allain and Samaha, 2019). Rats self-administered cocaine on a PR schedule for 4 days at 4 different doses (0.063, 0.124, 0.25, and 0.75 mg/kg/infusion), with the doses counterbalanced except the largest dose was always on the final day. After each cocaine infusion (paired with the 10-second audiovisual cue and timeout), the number of lever presses required to obtain the next infusion increased on a logarithmic scale. Sessions ended after 5 hours or when 1 hour passed without the next infusion being obtained.

#### 2.5.6 Cocaine-seeking tests after intra-DLS drug infusion

Rats in experiment 2 with implanted with DLS guide cannulae (n=48; male n=24; female n=24) and were given a total of 4 cocaine-seeking tests 15 minutes after bilateral infusion of drug (10 μg *cis*-flupenthixol) or vehicle (ddH_2_O) through a 28-guage internal cannula extending 1 mm below the guide cannula connected to a syringe pump (Harvard Apparatus) and 10 μl Hamilton syringes. Internal cannulae were left in place for one minute following infusion. Cocaine-seeking tests were identical to self-administration sessions except they were 15 minutes long and cocaine was not delivered upon timeout initiation. Rats immediately entered their daily cocaine self-administration session after each cocaine-seeking test. Tests occurred on slightly different timelines for two groups (Figure 3A, 3C). In both groups (n=24 each), tests 1-2 (after *cis*-flupenthixol or vehicle, counter-balanced) occurred prior to any IntA, and tests 3-4 occurred after IntA, allowing for a within-subjects analysis between drug and vehicle and before and after IntA. The second group was added to ensure 10 days of undisrupted IntA before tests 3-4.

**Figure 1:**
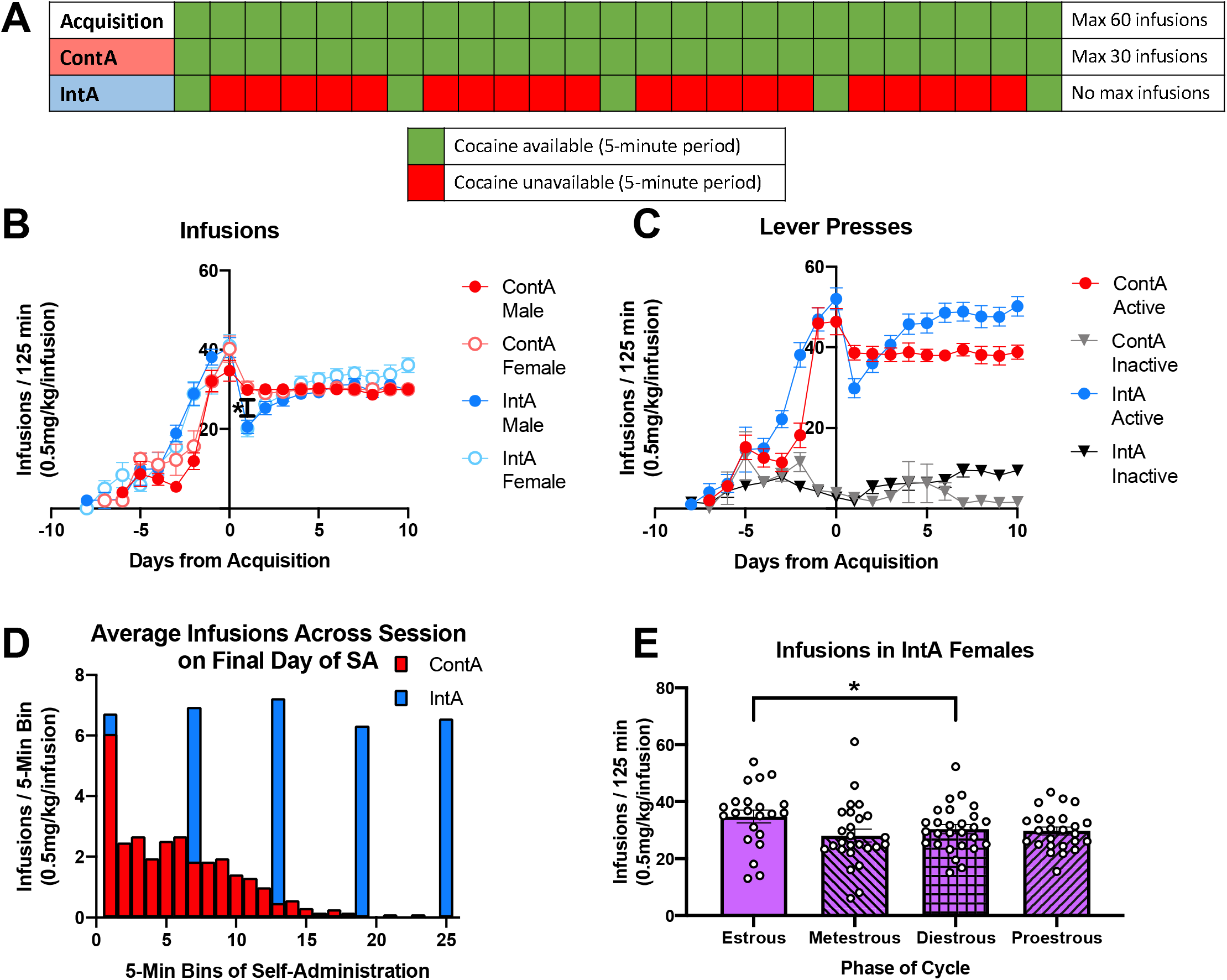
Continuous vs. intermittent cocaine self-administration produce different patterns of cocaine intake. Schematic for acquisition, ContA, and IntA self-administration sessions (A). All rats (experiments 1-2) self-administered cocaine for 2-10 days until they met acquisition criteria, then for 10 days with either ContA or IntA to cocaine. During the 10 days of ContA or IntA after acquisition, there was no effect of access model or sex on infusions, but there was a main effect of training day and a training day × access model interaction (B). Post-hoc analyses revealed that on the first day of ContA or IntA training, IntA-trained males and females self-administered less cocaine than ContA-trained males and females despite no overall differences in cocaine intake (B). For active and inactive lever presses during the 10 days of ContA or IntA after acquisition, there was a main effect of lever, training day, and access model, and there were significant training × access model, training × lever, and training × access model × lever interactions, but no access model × lever interaction (C). In a subset of rats, the number of infusions during each 5-minute period of the final day of self-administration were plotted in a histogram to show the different patterns of cocaine intake for ContA vs. IntA access models (D). The estrous phase in all female rats was monitored via vaginal cytology throughout self-administration. Although ContA almost always reached the maximum 30 infusions, there was a main effect of estrous phase on the average number of cocaine infusions in IntA-trained rats (E). Post-hoc analyses revealed that IntA-trained rats took more cocaine infusions in estrous than in diestrous (E). Graphs show group means ± SEM and individual data points. *p<0.05.

**Figure 2:**
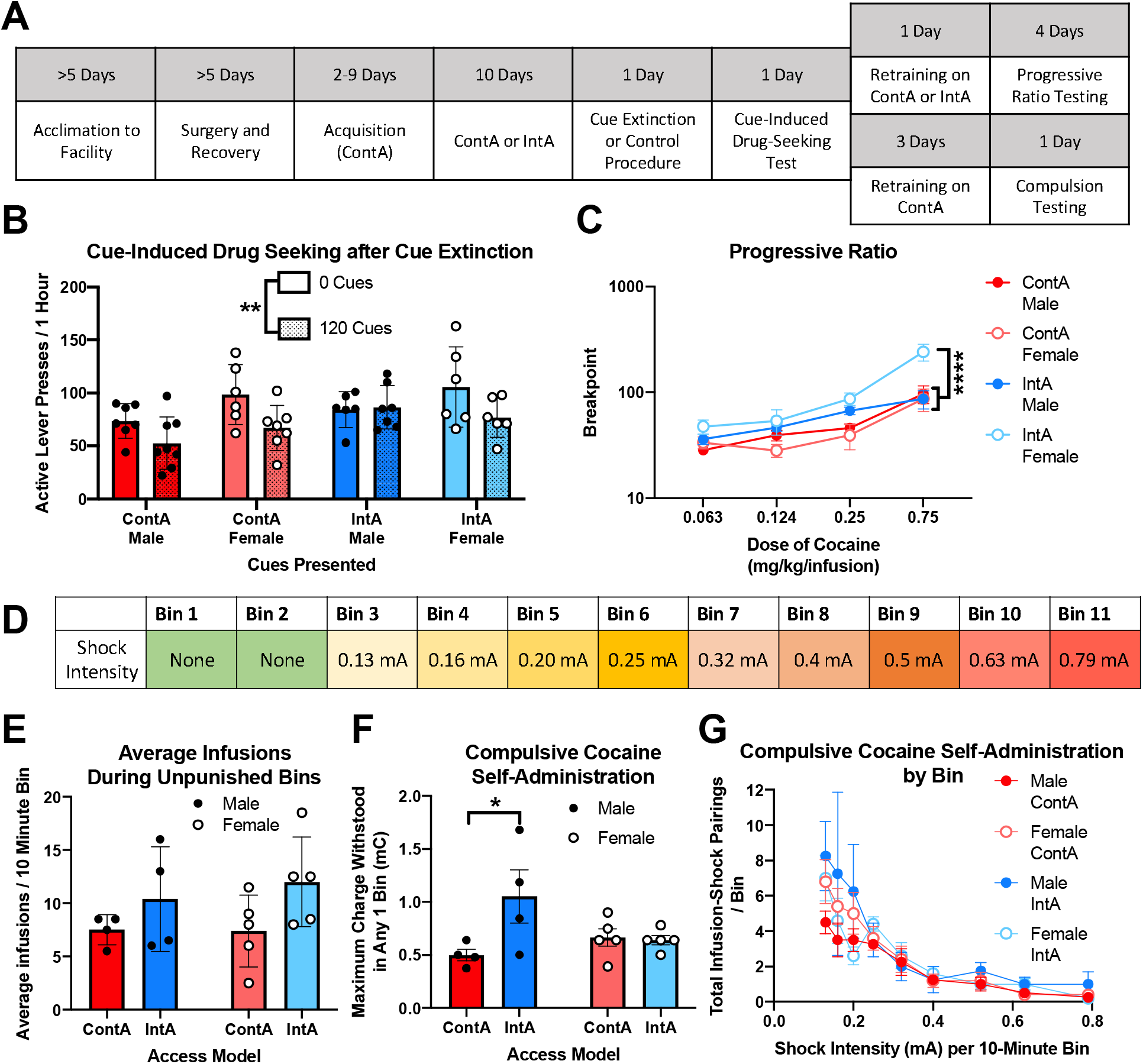
Sex-specific effects of intermittent cocaine self-administration on the efficacy of cue extinction, motivation for cocaine, and compulsive cocaine self-administration. Experimental timeline for the subset of rats included in experiment 1 (A). After 10 days of ContA or IntA, rats underwent cue extinction (120 CS) or a control procedure (0 CS) followed by a 1-hour cue-induced cocaine-seeking test. There was a main effect of cue extinction and access model on active lever presses, and overall rats that underwent 120-CS cue extinction made significantly fewer lever presses than 0 CS control rats (B). A subset of rats underwent progressive ratio testing for motivation for cocaine. There was a main effect of dose of cocaine, access model, and sex on the breakpoint in the progressive ratio test, and there were dose × access model, dose × sex, access model × sex, and dose × access model × sex interactions (C). Post-hoc analyses revealed that at the highest dose, IntA-trained females had a higher breakpoint than all other groups (C). Another subset of rats underwent a test for compulsive cocaine self-administration that involves a period of unpunished cocaine self-administration, followed by 10-minute bins where cocaine infusions are paired with a footshock of increasing intensity (D). There was no effect of sex or access model on the average number of cocaine infusions taken during unpunished bins (E). When the maximum footshock charge withstood in any 1 bin for each rat was calculated, there was an access model × sex interaction (F). Post-hoc analyses revealed that IntA-trained males withstood significantly more shock than ContA-trained males, but this was not the case for IntA-trained females vs. ContA-trained females (F). There was a main effect of shock intensity on the number of cocaine infusion-shock pairings per bin (G). Graphs show group means ± SEM and individual data points. *p<0.05. **p<0.01. ****p<0.0001.

**Figure 3:**
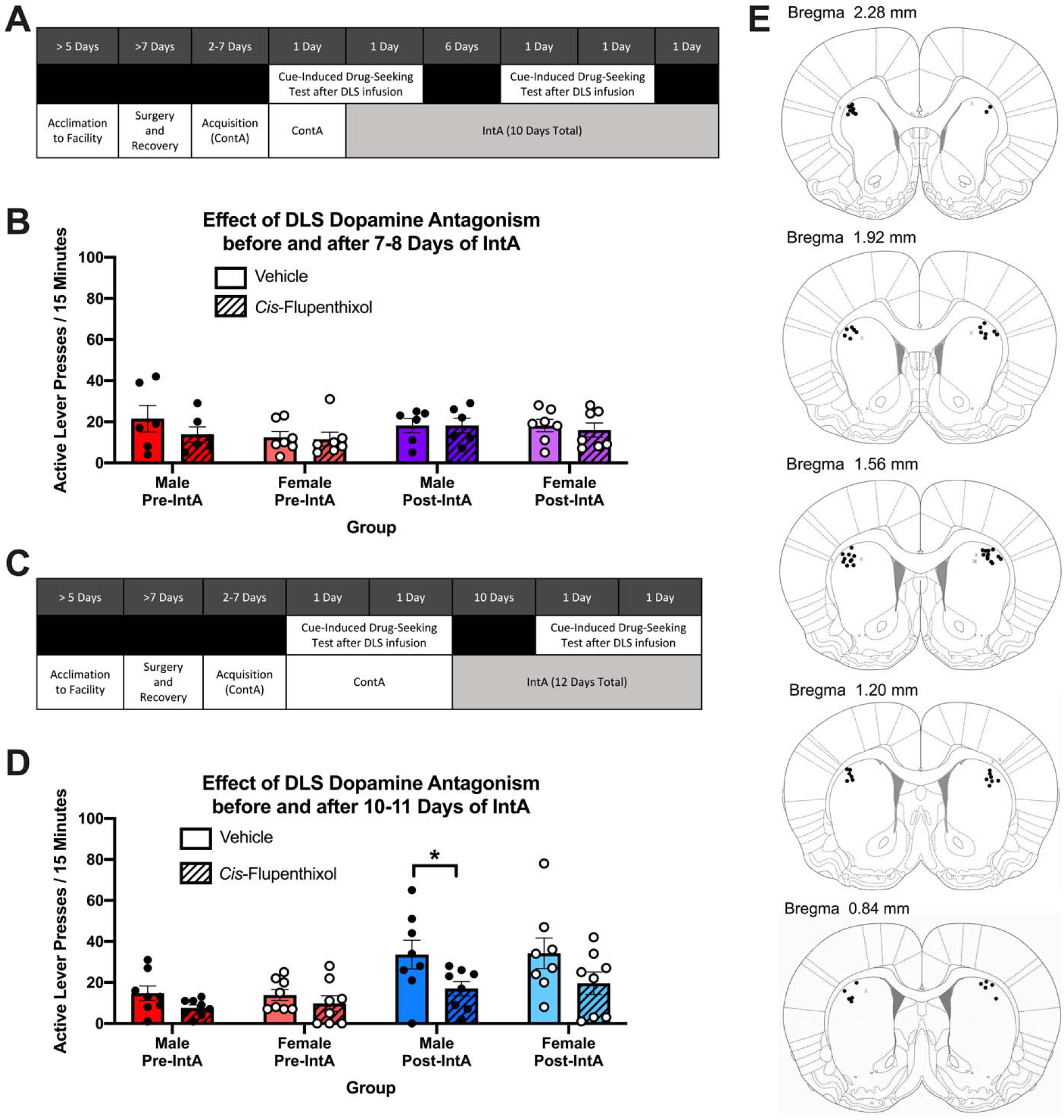
Intermittent cocaine self-administration facilitates cocaine seeking dependent on DLS dopamine. Experimental timeline for the subset of rats in experiment 2A (A). After acquisition (before IntA) and after 7-8 days of IntA, rats underwent a 15-minute drug-seeking test after direct DLS infusion of the nonspecific dopamine antagonist *cis*-flupenthixol or vehicle. There was no effect of DLS dopamine antagonism on drug seeking before or after 7-8 days of IntA (B). Experimental timeline for the subset of rats in experiment 2B (C). There was a main effect of drug and IntA training, as well as a drug × IntA training interaction on active lever presses during the cocaine-seeking tests before and after 10-11 days of IntA (D). Post hoc analyses revealed that males had fewer lever presses after DLS-dopamine antagonism with *cis*-flupenthixol compared to vehicle after, but not before, 10-11 days of IntA training (D). Placement of DLS cannulae were histologically evaluated, and each on-target placement is indicated by a black dot, while off-target misses are indicated by gray Xs (E). Graphs show group means ± SEM and individual data points. *p<0.05.

#### 2.6 Vaginal Cytology

After daily self-administration, vaginal cytology was used to monitor the estrous cycle of female rats. Sterile saline (200 μl) was pipetted into the vaginal canal and then placed onto a slide and coverslipped. Males were handled similarly. Slides were visualized at 200× magnification to determine estrous phase (Goldman et al., 2007).

#### 2.7 Histology

Rats in experiment 2 were euthanized with CO_2_ followed by decapitation, and brains were removed and submerged for at least 24 hours in 10% buffered formalin phosphate (Fischer Chemical). Brains were then cryo-protected with a 30% sucrose solution prior to sectioning and visualization as previously described (Bender and Torregrossa, 2021). Histological misses were characterized by placement more than 2.3 mm anterior to Bregma, less than 0.6 mm anterior to Bregma, dorsal of the striatum, or medial or lateral to the DLS.

#### 2.8 Exclusion Criteria

Rats were excluded from analysis due to death or illness after surgery (n=4), failure to meet acquisition criteria by 10 days (n=3), loss of catheter patency (determined by a 0.1-0.2 ml intravenous infusion of 10 mg/ml sodium brevital) (n=19), or if they failed to maintain an overall average of 15 daily infusions (n=3). Some rats were included in initial analysis and excluded from later analysis if catheter patency was lost between testing (n=5). For rats with histological misses (n=9), only self-administration data was included. Numbers of subjects reported in the results indicate the number of rats included in analysis after exclusions.

#### 2.9 Quantification and Statistical Analysis

Animals were matched for responding when split into groups, and experimenters were blind to rats’ treatment condition when possible. Behavioral data were collected using MedPC Software. All statistical analyses were performed on GraphPadPrism and SPSS Statistics software. The Shapiro-Wilk test was used to determine all data were normally distributed, and the Bartlett’s test was used to determine that there were no significant differences between groups in estimated variance. For all statistical analyses, significance was set at p<0.05. Infusions and lever presses during the 10 days of ContA or IntA were analyzed using a three-way rmANOVA with time as the repeated measure factor and either access model, sex, or lever as factors where indicated. All other analyses used two- and three-way ANOVAs as necessary, except a mixed-effects model was used to analyze the effect of estrous phase on infusions because not all rats exhibited every phase. When interactions were significant in ANOVA or mixed-effects analysis, Sidak’s or Tukey’s post-hoc multiple comparisons were used to further analyze differences between groups.

## 3. Results

### 3.1 Continuous vs. intermittent cocaine self-administration produce different patterns of cocaine intake

Self-administration data for all rats used in these studies are reported together (Figure 1). Rats were given continuous access to cocaine until acquisition criteria were met (days −10-0), and then were split into ContA and IntA groups for an additional 10 days of self-administration (days 1-10). During acquisition and ContA, rats could continuously self-administer cocaine until the maximum number of infusions were met, but during IntA cocaine availability was restricted to 5 5-minute periods (Figure 1A).

Rats (n=91) learned to self-administer cocaine, and during the 10 days after acquisition, there was no main effect of access model (F_(1,87)_=0.04217, p=0.8378) or sex (F_(1,87)_=0.9945, p=0.3214), but there was a main effect of training day (F_(9,783)_=9.860, ****p<0.0001) and a training day × access model interaction (F_(9,783)_=9.834, ****p<0.0001) (3-way rmANOVA) (Figure 1B). Further post-hoc analyses (Tukey’s multiple comparisons) indicated this interaction was driven by fewer infusions in the IntA-trained rats compared to the ContA-trained rats on the first day of IntA or ContA. IntA-trained males took significantly less cocaine than ContA-trained males (*p=0.0484, q=5.531) and ContA-trained females (*p=0.0374, q=5.635), and IntA-trained females took significantly less cocaine than ContA-trained males (*p=0.0407, q=5.601) and ContA-trained females (*p=0.0311, q=5.708). ContA-trained males and females (p>0.9999, q=0.3256) and IntA-trained males and females (p>0.9999, q=0.3363) did not differ in cocaine intake on the first day of IntA or ContA (Figure 1B). This reduction in infusions on the first day of IntA is likely due to the change in access model, when rats must learn they can only take cocaine in 5-minute periods. Note that we chose to cap ContA-trained rats at a maximum of 30 infusions to ensure that ContA- and IntA-trained rats were obtaining an equivalent number of cocaine infusions. Therefore, any differences in later tests would be due to the access model and would not be confounded by overall cocaine intake. Most ContA-trained rats reached the maximum within about an hour.

For lever presses during the 10 days after acquisition, there was a main effect of training day (F_(2.764,246)_=6.636, ***p=0.0004), access model (F_(1,89)_=5.958, *p=0.0016), and lever (F_(1,89)_=597.4, ****p<0.0001) (3-way rmANOVA) (Figure 1C). Additionally, there was a significant training × access model interaction (F_(9,801)_=6.777, ****p<0.0001), training day × lever interaction (F_(5.727,509.7)_=2.174, *p=0.0470), training day × access model × lever interaction (F_(9,801)_=3.174, ***p<0.0009), but no access model × lever interaction (F_(1,89)_=0.3738, p=0.5425). These data suggest that active lever presses increased as training progressed in IntA-trained rats, again likely due to the change in access model leading to a reduction in lever presses on the first few days of IntA until binge-like cocaine intake was learned. Despite having equivalent cocaine intake to ContA-trained rats, IntA-trained rats made more active lever presses by the end of training. This is likely due to increased timeout active lever presses, which occur during the timeout period and thus are counted but do not result in cocaine infusions. Although ContA and IntA-trained rats self-administered equivalent cocaine infusions, their patterns of cocaine intake were unsurprisingly very different. The average number of cocaine infusions during each 5-minute bin on the final day of self-administration was plotted for a subset of rats (n=37) (Figure 1D). ContA-trained rats binged in the beginning of the session and then reduced their cocaine intake to maintain their desired brain cocaine concentration until they reached the maximum 30 infusions, whereas IntA-trained rats binged during each 5-minute period of cocaine availability.

Vaginal cytology was used to monitor estrous phase in female rats throughout the study. Although female ContA-trained rats almost always reached the maximum 30 infusions, so no change could be observed, we examined if there was an effect of estrous cycle on the number of cocaine infusions obtained in IntA-trained females that completed 10 days of IntA (n=28). The average number of daily infusions each rat obtained during each estrous phase was calculated. There was a main effect of estrous phase on average number of cocaine infusions (F_(2.070,47.61)_=3.272, *p<0.0450) (mixed-effects analysis) (Figure 1E). Further post-hoc analyses (Tukey’s multiple comparisons) indicated rats took more infusions during estrous than diestrous (*p=0.0439, q=4.047) (Figure 1E). There were no other significant differences between phases, but there were nonsignificant trends towards rats taking more infusions in estrous than metestrous (p=0.0633, q=3.793) or proestrous (p=0.0588, q=3.862) (Figure 1E). Therefore, although IntA-trained females did not self-administer significantly more cocaine than IntA-trained males, they tended to take more cocaine during estrous than other cycle phases, though this effect was small.

### 3.2 Intermittent cocaine self-administration has sex-specific effects on the efficacy of cue extinction, motivation for cocaine, and compulsive cocaine self-administration

After 10 days of ContA or IntA, rats in experiment 1 (n=53) underwent behavioral testing for the effects of cue extinction on cue-induced drug seeking (n=53), motivation for cocaine using a PR test (n=30), and compulsive cocaine self-administration (n=18) (Figure 2A). Because we have previously shown that rats trained on second-order schedules of reinforcement are resistant to cue extinction (Bender and Torregrossa, 2021), we first sought to determine if a different pattern of self-administration (such as IntA) could also facilitate this resistance to cue extinction. After the 10^th^ day of ContA or IntA, rats underwent cue extinction, when 120 audiovisual cues (120 CS) were presented non-contingently, or a control 0-CS procedure. The following day, rats underwent a 1-hour cue-induced cocaine-seeking test, and lever presses were recorded and resulted in cues but no cocaine delivery. There was a main effect of cue extinction (F_(1,45)_=8.995, **p<0.0044) and access model (F_(1,45)_=5.431, *p<0.0243) on active lever presses during the cue-induced cocaine-seeking test, but no main effect of sex (F_(1,45)_=3.680, p=0.0614), cue extinction × access model interaction (F_(1,45)_=0.9398, p=0.3375), cue extinction × sex interaction (F_(1,45)_=2.459, p=0.1239), access model × sex interaction (F_(1,45)_=1.108, p=0.2982), or 3-way interaction (F_(1,45)_=0.6079, p=0.4397) (3-way ANOVA) (Figure 2B). These results suggest that overall, cue extinction reduced cue-induced cocaine seeking, and IntA-trained rats showed increased cue-induced cocaine seeking than ContA-trained rats. Moreover, while the interaction effects did not reach statistical significance, visual inspection of the data suggests that the IntA-trained males, but not females, were resistant to the effects of cue extinction.

A subset of rats from experiment 1 (n=30) underwent PR testing to determine if there was an effect of IntA or sex on motivation for cocaine. For 4 days on 4 different doses of cocaine, the number of lever presses required to obtain the next cocaine infusion increased on a logarithmic scale. The ratio at which rats no longer obtained another infusion, or the breakpoint, is an indicator of motivation (Panlilio and Goldberg, 2007). There was a main effect of dose, (F_(3,78)_=41.07, ****p<0.0001), access model (F_(1,26)_=19.83, ***p=0.0001), and sex (F_(1,26)_=8.007, **p=0.0086) on breakpoint (3-way rmANOVA) (Figure 2C). There were also dose × access model (F_(3,78)_=4.435, **p=0.0062), dose × sex (F_(3,78)_=6.850, ***p=0.0004), access model × sex (F_(1,26)_=12.99, **p=0.0013), and 3-way dose × access model × sex interactions (F_(3,78)_=7.952, ***p=0.0001). Further post-hoc analyses (Sidak’s multiple comparisons) indicated that IntA-trained females had a higher breakpoint at the highest dose (0.75 mg/kg/infusion) than ContA-trained males (****p<0.0001 t=7.253), ContA-trained females (****p<0.0001 t=7.517), or IntA-trained males (****p<0.0001 t=7.956). These data indicate that IntA promoted increased motivation for 0.75 mg/kg/infusion cocaine exclusively in females.

Another subset of rats from experiment 1 (n=18) were used to determine if IntA promotes increased compulsive cocaine self-administration. We used a previously-established test where the session is divided into 10-minute bins, and for the first 2 bins rats self-administer unpunished to reach and begin to maintain their desired brain-cocaine concentration (Bentzley et al., 2014). For the remaining 9 bins, the intensity of an aversive footshock paired with cocaine infusions is increased incrementally from 0.13 mA to 0.79 mA (Figure 2D).

First, we examined if there were any differences in the average number of unpunished cocaine infusions during the first 2 bins between sexes or access models. There were no main effects of access model (F_(1,14)_=4.448, p=0.0534) or sex (F _(1,14)_=0.1852, p=0.6735) or interaction (F_(1,14)_=0.2369, p=0.6735) (2-way ANOVA) (Figure 2E), suggesting that there were no significant differences in unpunished cocaine self-administration. Even so, we noted that both IntA-trained males and females did take non-significantly more cocaine unpunished. Therefore, to ensure differences in compulsive cocaine self-administration were not due to differences in the amount of cocaine self-administered, we used the previously-established method of calculating the maximum charge withstood in each 10-minute bin of punished cocaine self-administration and comparing the maximum charge each rat withstood in any one bin. This value is similar to a breakpoint and represents the amount of punishment each rat is willing to withstand to maintain their desired brain-cocaine concentration. We found no main effect of access model (F_(1,14)_=4.483, p=0.0526) or sex (F_(1,14)_=0.9766, p=0.3398) on the maximum charge withstood in any 1 bin, but there was an access model × sex interaction (F_(1,14)_=5.332, *p=0.0367) (2-way ANOVA) (Figure 2F). Further post-hoc analyses (Sidak’s multiple comparisons) indicated that IntA-trained males withstood significantly more shock than ContA-trained males (*p=0.0202, t=2.969), but IntA-trained females did not withstand more shock than ContA-trained females (p=0.9874, t=0.1439). Additionally, we examined the total cocaine infusion-shock pairings in each bin. There was a main effect of shock intensity (F_(2.077,29.08)_=16.13, ****p<0.0001) on punished infusions, indicating rats reduced their infusions as the shock intensity increased (3-way rmANOVA) (Figure 2G). There were no main effects of access model (F_(1,14)_=2.742, p=0.1200) or sex (F_(1,14)_=0.01683, p=0.8986), nor any shock intensity × access model (F_(8,112)_=0.3812, p=0.9287), shock intensity × sex (F_(8,112)_=0.2420, p=0.9819), access model × sex (F_(1,14)_=4.339, p=0.0561), or 3-way (F_(8,112)_=1.099, p=0.3691) interactions. Overall, these results suggest that IntA enhanced compulsive cocaine self-administration exclusively in males.

### 3.3 Intermittent cocaine self-administration facilitates cocaine seeking dependent on DLS dopamine

Given that we have previously shown that DLS dopamine-dependent, habit-like cocaine seeking is resistant to cue extinction, the observation that IntA-trained males may be resistant to the effects of cue extinction led us to examine if DLS dopamine antagonism impacts cocaine seeking after IntA in experiment 2 (n=29). Rats were infused bilaterally in the DLS with the nonspecific dopamine antagonist *cis*-flupenthixol and vehicle (counterbalanced across 2 days) prior to a 15-minute drug-seeking test at two different timepoints before and after IntA for a total of 4 tests.

Initially, in experiment 2A (n=13), the first set of tests occurred after acquisition and prior to IntA, and the second set occurred after 7-8 days of IntA (Figure 3A). All rats received both drug and vehicle and were tested at both timepoints for a within-subjects design. For active lever presses during the cocaine-seeking tests after vehicle or *cis*-flupenthixol infusion, there was no effect of drug (F_(1,11)_=0.7622, p=0.4013), IntA (F_(1,11)_=1.248, p=0.2877), or sex (F_(1,11)_=1.282, p=0.2815), and there were no interactions between drug × IntA (F_(1,11)_=0.6104, p=0.4498), drug × sex (F_(1,11)_=0.8404, p=0.3789), IntA × sex (F_(1,11)_=0.8404, p=0.3789), or 3-way interaction (F_(1,11)_=1.209, p=0.2951) (3-way ANOVA) (Figure 3B). These data suggest that both before and after minimal (7-8) days of IntA, drug seeking is unaffected by DLS dopamine antagonism.

Experiment 2B (n=16) was very similar, but we made some adjustments to the timeline to ensure that no drug-seeking test occurred on the first day of IntA and rats would receive 10 days of undisrupted IntA prior to the second set of drug-seeking tests after DLS dopamine antagonism (Figure 3C). There was a main effect of drug (F_(1,14)_=16.75, **p=0.0011) and IntA (F_(1,14)_=15.73, **p=0.0014) on active lever presses during cocaine-seeking tests, but no effect of sex (F_(1,14)_=0.06260, p=0.8061) (3-way ANOVA) (Figure 3D). There was also a drug × IntA interaction (F_(1,14)_=4.667, *p=0.0486), but no drug × sex (F_(1,14)_=0.2318, p=0.6376), IntA × sex (F_(1,14)_=0.01838, p=0.8941), or 3-way interactions (F_(1,14)_=0.01167, p=0.9155) (Figure 3D). Further post-hoc analyses (Sidak’s multiple comparisons) indicated that males made fewer lever presses after DLS-dopamine antagonism with *cis*-flupenthixol compared to vehicle after (*p=0.0255, t=3.380), but not before (p=0.8741, t=1.449), at least 10 days of IntA training (Figure 3D). These results suggest that at least 10 days of IntA facilitated DLS dopamine-dependent cocaine seeking, and this effect was more pronounced in males but did appear in females as well. Histological analysis of DLS cannulae placement are shown (Figure 3E).

## 4. Discussion

Here, we showed notable sex differences in the effects of IntA on addiction-like behavior. We found that compared to ContA, just 10 days of IntA sessions that were just over 2 hours long were enough to promote several addiction-like behaviors differentially in males and females. Recent studies have utilized these 2-4-hour long IntA sessions (Allain and Samaha, 2019; Cippitelli et al., 2022; Garcia et al., 2020; Kawa et al., 2019), shorter than the traditional 6-8 hours, which are more accessible to labs with limited operant space or time. Interestingly, one study in males suggested that shorter IntA sessions promote more individual differences, where a subset of rats showed escalated cocaine intake accompanied by increased cue-induced reinstatement and locomotor sensitization to cocaine (Garcia et al., 2020). Despite using a similar protocol, we did not find a subgroup of IntA-trained rats (male or female) expressing escalated cocaine intake, likely because the previous study saw escalation occur mainly on days 10-14, which we did not include (Garcia et al., 2020). Still, we extend these findings by showing short IntA sessions also promote sex differences in the effects of IntA, indicating that these protocols may be useful in identifying risk factors that contribute to acceleration of SUD development during early stages of drug use.

Although we found no sex differences in cocaine intake, IntA-trained females took more cocaine when in estrous, replicating previous findings in a ContA models (Feltenstein and See, 2007; Roberts et al., 1989). We could not determine if estrous phase had an effect on ContA-trained females in our study because they almost always obtained the maximum number of infusions, and therefore no differences could be observed. Additionally, our experiments were not powered to determine if estrous phase impacted the effects of IntA on the various aspects of addiction-like behavior we evaluated. Given our findings and previous studies showing that estradiol can influence sensitization to cocaine, which is enhanced in IntA-trained females, future experiments should determine the effect of circulating sex hormones on the ability of IntA to promote addiction-like behaviors in females (Carr et al., 2020; Sell et al., 2002).

We did not find an enhancement of motivation for cocaine in males after IntA compared to ContA, which was surprising given several studies that have shown this enhancement using either PR or behavioral economic testing (Algallal et al., 2020; Calipari et al., 2015; James et al., 2019; Zimmer et al., 2012). Even though evidence suggests that the length of daily IntA does not impact motivation (Allain and Samaha, 2019), because our IntA training was rather minimal (10 days) compared to these other studies, it is possible that more days of short IntA training are necessary to enhance motivation for cocaine in males. Alternatively, it is possible that motivation for cocaine in males was enhanced but was masked by an over-reliance on the DLS dopamine-dependent, habit-like behavior we identified in IntA-trained males. Both the PR and behavioral economic tests for motivation rely on the assumption that rats are responding in a goal-directed manner and increasing their lever presses when the outcome is of higher value, but if behavior is habit-like and does not rely on the value of the outcome, breakpoint may not be an accurate measure of motivation. Future studies should investigate if restoring goal-directed control could reveal increased motivation in a PR test in IntA-trained males. We did find that IntA-trained females showed increased motivation compared to IntA-trained males, which agrees with previous literature suggesting enhancement of motivation for cocaine after IntA is greater in females than in males (Algallal et al., 2020; Kawa and Robinson, 2019). Additionally, we found that IntA-trained females showed greater motivation for cocaine compared to ContA-trained females, as previously shown (Algallal et al., 2020). Based on other studies, this increased motivation in females may be reduced when females are in proestrous or increased during estrous (Kohtz et al., 2022; Roberts et al., 1989), but it is unlikely that circulating hormones in females contributed to this effect in our hands because estrous phases were fairly balanced between ContA and IntA groups during PR testing. Overall, these findings along with the previous literature suggest that IntA promotes increased motivation for cocaine more quickly and more robustly in females, so future studies should examine the role of sex hormones and biological sex differences on the interaction between sensitization and motivation for cocaine.

Our findings expand on previous research showing that IntA promotes increased compulsive cocaine self-administration in males (James et al., 2019), but we also showed a lack of this effect in females. These results suggest that IntA promotes increased compulsive cocaine self-administration either more quickly or exclusively in males, and future studies should determine if more days of IntA or longer daily IntA sessions would be sufficient to replicate this effect in females. We also determined that IntA-trained rats reduced active lever presses after DLS dopamine antagonism after a full 10-11 days of IntA, but not before, suggesting the development of habit-like, DLS dopamine-dependent behavior, most notably in male rats. This is a novel finding and should be examined further using behavioral analysis of habitual behavior to supplement the pharmacology used here, since no previous studies have examined how IntA may promote the development of habitual behavior. There is evidence that functional disconnection of the OFC and ventrolateral striatum, a circuit shown to be involved in goal-directed behavior, reduces responding on a PR schedule in IntA-trained rats (Gourley et al., 2013; Minogianis et al., 2019), which contradicts our findings by suggesting that self-administration under a PR schedule after IntA remains goal-directed. However, this experiment did not examine the DLS and only allowed rats to take up to 0.5 mg/kg/infusion during each drug-available period of training, which is well below what most rats would take freely in an IntA binge period (Minogianis et al., 2019). It is possible that a combination of IntA and higher doses of cocaine are required to facilitate DLS dopamine-dependent behavior, and this should be further investigated.

We also examined the ability of Pavlovian cue extinction to reduce drug seeking after ContA or IntA. We did show overall increased lever presses in IntA-trained compared to ContA-trained rats, in accordance with previous studies showing increased cue-induced drug seeking in IntA-trained rats (Aragona et al., 2009; James et al., 2019; Kawa et al., 2019). Additionally, we found a main effect of cue extinction to reduce lever pressing in a cue-induced drug-seeking test, providing new evidence that despite enhanced cue-mediated responding, drug seeking in IntA-trained rats is still susceptible to extinction of the drug-cue association. However, it should be noted that the cue extinction effect in IntA-trained rats was driven by the females and was not visually apparent in males, suggesting the study was underpowered to detect a 3-way interaction. Moreover, given that the IntA-trained males were also more affected by intra-DLS dopamine antagonism and showed more compulsive-like cocaine self-adminstration, it is possible that males trained on an IntA schedule have a greater susceptibility to developing habit-like cocaine seeking. This is further supported by our previous findings that DLS dopamine-dependent behavior is resistant to cue extinction after training on a second-order schedule (Bender and Torregrossa, 2021). Notably, second-order training produced this habit-like behavior in both males and females, suggesting that the amount of IntA training used in the present study may have revealed a slight sex difference in the propensity of cocaine seeking to become DLS dopamine-dependent, but that with more training females would likely exhibit habit-like behavior similar to males. This result is further supported by previous work suggesting that genetically male mice are more prone to form ethanol-seeking habits than females (Barker et al., 2010), though the opposite is true for the seeking of a food reward (Quinn et al., 2007).

Interestingly, because we found increased PR breakpoint exclusively in IntA-trained females, but increased compulsive behavior exclusively in IntA-trained males, our results suggest that compulsive cocaine self-administration and motivation for cocaine are promoted independently. Additionally, because the IntA-trained males showed the most robust DLS dopamine-dependent behavior, it may be that the development of habit-like behavior coincides with the degree of punishment-resistant cocaine seeking. Moreover, that IntA may promote habit-like behavior, increased motivation for cocaine, and compulsive behavior in a sex-specific manner lends credence to its usefulness in modeling addiction-like behavior in rodents and examining the different contributions of these drug effects on behavior. Importantly, the sex differences we identified in these experiments occurred after rather limited IntA. We may be capturing a timepoint at which these addiction-like behaviors may be developing in some rats at different rates based on sex and other individual differences. It is possible that additional short IntA training, such as 10-20 additional days, would promote enhanced motivation, compulsive cocaine self-administration, resistance to cue extinction, and reliance on DLS dopamine across sexes, and future experiments should examine this. Overall, we expanded on the current literature by providing new insights into how IntA may differentially promote addiction-like behaviors in males and females. Our findings suggest that IntA may be uniquely suited to identify sex differences in how cocaine impacts dopamine modulation, motivation, and punishment resistance, especially during the early stages of drug use when sensitization occurs.

## Author Contributions

BNB: Conceptualization, methodology, software, investigation, formal analysis, writing – original draft, writing – review & editing, project administration. MMT: Conceptualization, methodology, software, formal analysis, writing – review & editing, visualization, supervision, funding acquisition.

## Funding and Acknowledgements

The research was supported by the National Institute of Health grant R01DA042029 (M.M.T.). We would like to acknowledge Dana Smith, Camryn Forbes, Michael Wright, Alexis Egazarian, Lauren Charlton, Lindsey Buchman, Alina Owsiany, and Juan Robayo for assistance with behavioral experiments, surgical procedures, and histology, and Sierra Stringfield for technical advice.

